# Optimizing Tissue Lysis and DNA Extraction Protocols to Enhance Bacterial Diversity Profiling in the *Drosophila melanogaster* Gut Microbiome

**DOI:** 10.1101/2025.11.15.688634

**Authors:** Carlos L. Quiñones-Sanchez, Jan L. Bilbao-Del Valle, Miguel A. Urdaneta-Colon, Tasha M. Santiago-Rodriguez, Imilce A. Rodriguez-Fernandez

**Author notes:** These authors contributed equally to this work (Carlos L. Quiñones-Sanchez, Jan L. Bilbao-Del Valle).

## Abstract

The gut microbiota is a dynamic community that influences host metabolism, immunity, and overall health. Accurate characterization of this community requires robust and reproducible DNA extraction methods; however, technical biases introduced during tissue lysis and DNA isolation remain major challenges in microbiome research, particularly in animal model systems. In this study, we compared two commercial DNA extraction kits (Qiagen and Zymo) and two lysis methods (manual pestle homogenization and bead-beating) to evaluate their impact on microbiota profiling in a microbial community standard (MCS) and *Drosophila melanogaster* gut samples, a tractable model for host–microbe interactions. Full-length 16S rRNA sequencing was performed using Oxford Nanopore Technologies, followed by bioinformatic analysis using EPI2ME for taxonomic classification and standard diversity pipelines. Our data revealed that extraction and lysis methods significantly influence microbial composition, with some protocols resulting in inflated richness in MCS samples. Pestle homogenization with the Qiagen kit yielded the highest bacterial species richness while maintaining consistent representation of both Gram-positive and Gram-negative taxa. These findings demonstrate that extraction methodology strongly affects microbial diversity estimates and emphasize the need for standardized protocols to ensure reproducibility across microbiome studies, particularly those using model systems.

## Introduction

The microbiome, often referred to as an “unseen organ,” encompasses the diverse communities of bacteria, fungi, viruses, archaea, and eukaryotes that inhabit host organisms and interact with one another. These microorganisms play essential roles in maintaining physiological balance, regulating immunity, synthesizing metabolites and amino acids, and providing protection against pathogens[^1–5^]. Factors such as age, diet, infection, and antibiotic exposure can disrupt this balance, leading to microbiome dysbiosis and associated health disorders[^3,4^].

Advances in sequencing technologies have transformed microbiome research. Whole-genome sequencing (WGS) enables species-level resolution and functional profiling, while 16S rRNA gene sequencing (16S) remains a cost-effective method for taxonomic characterization[^5–7^]. Traditional short-read approaches typically target partial variable regions (e.g., V3–V4), limiting classification to the genus level. Full-length 16S sequencing (∼1,500 bp) overcomes this limitation by improving species-level resolution and accuracy[^6,8,9^].

Long-read sequencing technologies now permit near full-length 16S amplicon sequencing. Oxford Nanopore Technologies (ONT) and PacBio platforms generate reads that span the entire 16S gene, enhancing taxonomic precision. ONT’s portable and low-cost MinION sequencer provides an accessible option for laboratories without large-scale infrastructure, whereas PacBio’s HiFi sequencing offers higher raw accuracy but at greater cost[^10–14^]. A side-by-side comparison demonstrated that PacBio unique molecular identifier (UMI) sequencing achieved the highest accuracy (∼0.0007% error rate), while ONT UMI sequencing provided a lower-cost, field-deployable alternative[^15^]. Despite ONT’s higher per-read error rate (∼1%), continuous improvements in flow-cell chemistry and basecalling algorithms are steadily narrowing this accuracy gap[^11^].

ONT also offers integrated, user-friendly solutions such as the EPI2ME cloud platform for demultiplexing, taxonomic assignment, and diversity analysis, allowing users with limited computational experience to process data without high-performance computing resources and programming expertise[^16^]. This accessibility makes ONT particularly suitable for microbiome studies in small or resource-limited laboratories.

Accurate microbiome profiling depends not only on sequencing technology but also on effective sample lysis and DNA extraction. The efficiency of cell disruption during these steps is critical as microbial cells in complex materials, such as host tissues, can be difficult to lyse. Suboptimal extraction can bias species detection, particularly for Gram-positive bacteria with thick peptidoglycan walls[^17–21^]. Methodological differences at this step can therefore influence downstream diversity metrics and taxonomic profiles.

Although most microbiome studies focus on humans or murine models, *Drosophila melanogaster* has emerged as a powerful system for dissecting host–microbiome interactions[^22–24^]. The fly gut microbiota is simple, dominated by genera such as *Acetobacter* (Gram-negative), and *Lactobacillus* (Gram-positive), allowing precise manipulation and controlled experiments. Its tractable genetics and short life cycle make it an ideal model for testing how methodological workflows and interventions influence the gut microbiome composition and abundance.

In this study, we compared four DNA extraction protocols combining two commercial kits and two tissue homogenization methods to determine which yields the most comprehensive and reproducible bacterial profiles in *Drosophila* gut samples and a commercially available microbial community standard (MCS). By integrating alpha and beta diversity analyses, taxonomic composition, and Gram-stain classification, we aimed to identify the extraction protocol that maximizes bacterial detection while maintaining accuracy and reproducibility for *Drosophila* gut samples. This optimized workflow provides a practical framework for reliable, Nanopore-based microbiome analyses in low-resource laboratory environments without access to high-performance computing facilities, and underscores the importance of protocol standardization to ensure consistency and comparability across studies.

## Results

### Taxonomic classification of the mock community revealed inflated species-level assignments

We compared the performance of two commercially available DNA extraction kits and two tissue homogenization methods in recovering representative bacterial DNA from *D. melanogaster* gut samples (**Table 1**). The four protocols - BPQK, BPZK, ZBQK, and ZBZK - represent combinations of Blue Pestle (BP) or Zymo Bead Beating (ZB) homogenization with the Qiagen DNeasy Blood & Tissue Kit (QK) or the Zymo DNA Miniprep Kit (ZK), respectively. To evaluate the extraction performance, a microbial community standard (MCS) consisting of five Gram-positive, three Gram-negative bacterial and two yeast species in equal abundance was included as a control. The MCS underwent the same tissue lysis and genomic DNA (gDNA) extraction procedures as the *Drosophila* samples. Full-length 16S sequencing identified the expected bacterial genera across all four extraction protocols (**Figure 1A**). However, species-level classification revealed inflation of taxonomic assignments (herein referred to as “Inflated data”), with several community members misidentified or overrepresented (**Figure 1B**). This inflation affected the inferred Gram-stain distribution, leading to an apparent dominance of Gram-positive taxa across all conditions (**Figure 1C**).

**Figure 1.**
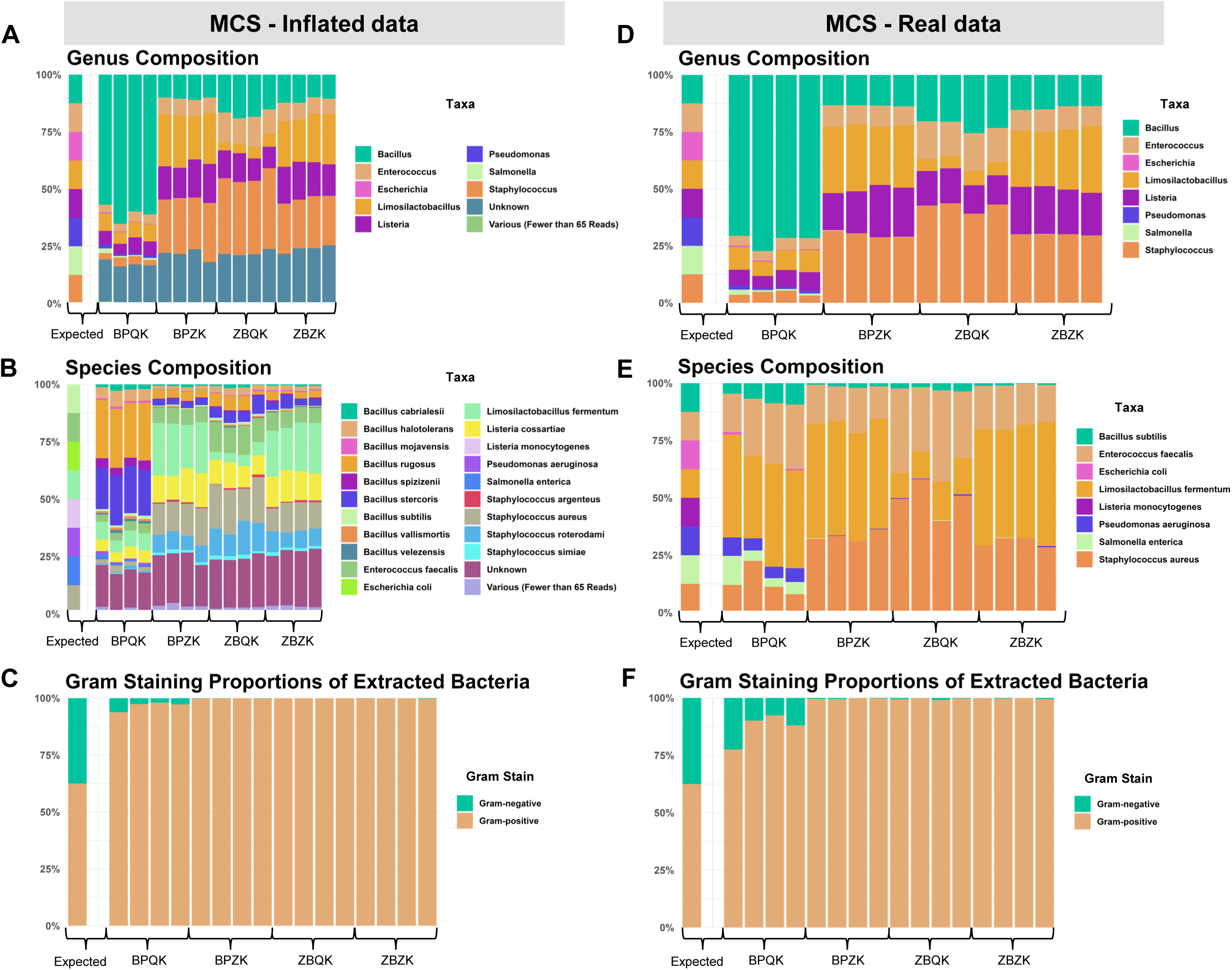
Taxonomic composition and Gram-stain distribution of the microbial community standard (MCS) across four DNA extraction protocols. **(A–C)** *Inflated data.* Full-length 16S rRNA sequencing of the MCS after DNA extraction using four protocol combinations (BPQK, BPZK, ZBQK, and ZBZK), representing Blue Pestle (BP) or Zymo Bead Beating (ZB) homogenization paired with the Qiagen DNeasy Blood & Tissue Kit (QK) or Zymo DNA Miniprep Kit (ZK). **(A)** Genus-level composition showing the expected and observed distributions. **(B)** Species-level composition revealing inflation of taxonomic assignments. **(C)** Gram-stain proportions inferred from species assignments, showing an apparent dominance of Gram-positive taxa. (**D–F)** *Real data.* Taxonomic composition after bioinformatic filtering to remove misidentified taxa and retain only true positives. **(D)** Genus-level composition of the filtered dataset. **(E)** Species-level composition representing the true mock community members. **(F)** Gram-stain proportions of the filtered dataset, showing that only the BPQK protocol recovered a noticeable fraction of DNA from Gram-negative bacteria. *Abbreviations:* BPQK = Blue Pestle + Qiagen Kit; BPZK = Blue Pestle + Zymo Kit; ZBQK = Bead Beating + Qiagen Kit; ZBZK = Bead Beating + Zymo Kit.

**Table 1.** Overview of DNA extraction protocols used in this study.

| Sample ID | Sample Type | Replicates | Homogenization Method | Extraction Method | Sequencing Run |
| --- | --- | --- | --- | --- | --- |
| BPQK | Microbial Community Standard | 4 | Blue Pestle | Qiagen DNeasy Blood&Tissue Kit | Run 1 |
|  | *Drosophila* | 3 (10 guts per replicate) |  |  |  |
| ZBQK | Microbial Community Standard | 4 | Bead Beating | Zymo DNA Miniprep Kit | Run 1 |
|  | *Drosophila* | 3 (10 guts per replicate) |  |  |  |
| BPZK | Microbial Community Standard | 4 | Blue Pestle | Qiagen DNeasy Blood&Tissue Kit | Run 2 |
|  | *Drosophila* | 3 (10 guts per replicate) |  |  |  |
| ZBZK | Microbial Community Standard | 4 | Bead Beating | Zymo DNA Miniprep Kit | Run 2 |
|  | *Drosophila* | 3 (10 guts per replicate) |  |  |  |

After bioinformatically removing misidentified taxa and retaining only true positives (herein referred to as “Real data”), we evaluated each extraction protocol by comparing the observed taxa with the expected composition of the MCS. At the genus level, removing unexpected taxa had little effect on the overall composition, which remained similar to that observed in the inflated dataset (**Figure 1D**). For instance, in all BPQK samples, the genus *Bacillus* dominated (∼70%), whereas other genera were detected at lower relative abundances. In contrast, *Bacillus* was less abundant in the remaining protocols, which showed higher proportions of *Staphylococcus, Listeria,* and *Limosilactobacillus*. Species-level filtering yielded the real composition of the mock community (**Figure 1E**), with *Limosilactobacillus fermentum* being highly abundant across all samples and *Staphylococcus aureus* prevalent in BPZK, ZBQK, and ZBZK samples. Filtering also modified the Gram-stain proportions (**Figure 1F**); among all protocols, only BPQK recovered a noticeable fraction of DNA from Gram-negative bacteria.

To further evaluate the taxonomic accuracy of each extraction protocol, true positives, false positives, and false negatives were quantified and visualized using Venn diagrams, at the genus level (**Figure 2**) and at the species level (**Supplementary Figure 1; Supplementary Data 2**). In BPQK samples, all eight expected bacterial genera were correctly identified, though four false positives (*Citrobacter*, *Shigella*, *Mammaliicoccus* and several unknown) were also detected (**Figure 2A**). Both BPZK and ZBQK protocols recovered most expected genera, with *Escherichia* being the only false negative (**Figures 2B and 2C**). These protocols also showed the presence of *Mammaliicoccus* and *Macrococcus* as false positives. In contrast, the ZBZK protocol failed to detect both *Salmonella* and *Escherichia*, but no false positives were observed in these samples, although some unknowns were present (**Figure 2D**).

**Figure 2.**
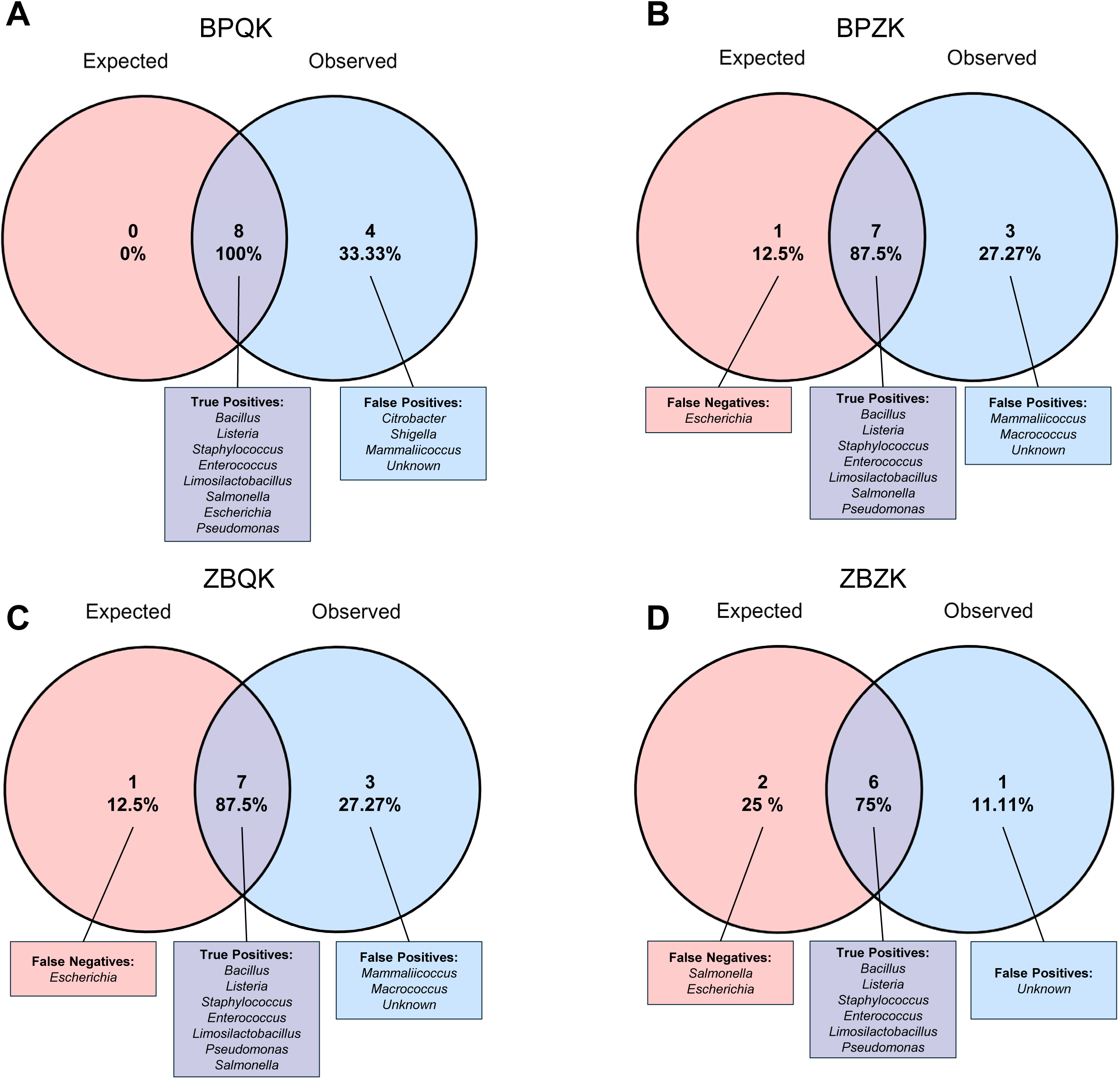
Comparison of expected and observed bacterial genera in the microbial community standard (MCS) across four DNA extraction protocols. Each Venn diagram illustrates the overlap between expected and observed bacterial genera detected after full-length 16S rRNA sequencing of the MCS. The left circle represents the eight genera expected in the mock community, while the right circle represents the genera detected for each extraction protocol. Percentages indicate the proportions of shared taxa (true positives), undetected expected taxa (false negatives), and taxa not present in the mock community (false positives). (**A**) BPQK (Blue Pestle + Qiagen Kit), (**B**) BPZK (Blue Pestle + Zymo Kit), (**C**) ZBQK (Bead Beating + Qiagen Kit), and (**D**) ZBZK (Bead Beating + Zymo Kit).

At the species level, all four protocols detected over 30 false positives in their samples (**Supplementary Figure 1**). The BPQK method detected 7 of the 8 expected species, representing the highest number of true positives among all extraction protocols (**Supplementary Figure 1A**), whereas the remaining protocols detected only 5–6 of the expected species (**Supplementary Figure 1B-1D**). To evaluate the relationships between false-positives and true-positive taxa, we calculated the p-distances between the reference sequences used by EPI2ME for taxonomic classification. These relationships were visualized using a Neighbor-Joining Phylogenetic Tree constructed in MEGA12 (**Supplementary Figure 2**). Most false positives clustered within the same genera as their corresponding true positives, with pairwise sequence identities exceeding 99% in most cases.

### Inflation of mock community data led to biases in alpha and beta diversity

To determine how data inflation affected diversity estimates, we compared the observed species richness, Shannon diversity index, and Bray-Curtis dissimilarities between the MCS inflated and filtered (real) MCS datasets relative to the expected community (**Figure 3**). Before filtering, all extraction protocols showed a trend toward higher alpha diversity values than expected, as reflected by both observed species richness (**Figure 3A**) and Shannon diversity index (**Figure 3B**). However, differences in alpha diversity among extraction protocols were not statistically significant (Kruskal–Wallis, p > 0.05). In contrast, Bray-Curtis dissimilarities revealed significant differences among protocols and the expected community, with BPQK samples forming a distinct cluster separate from the other methods and from the expected composition (**Figure 3C**).

**Figure 3.**
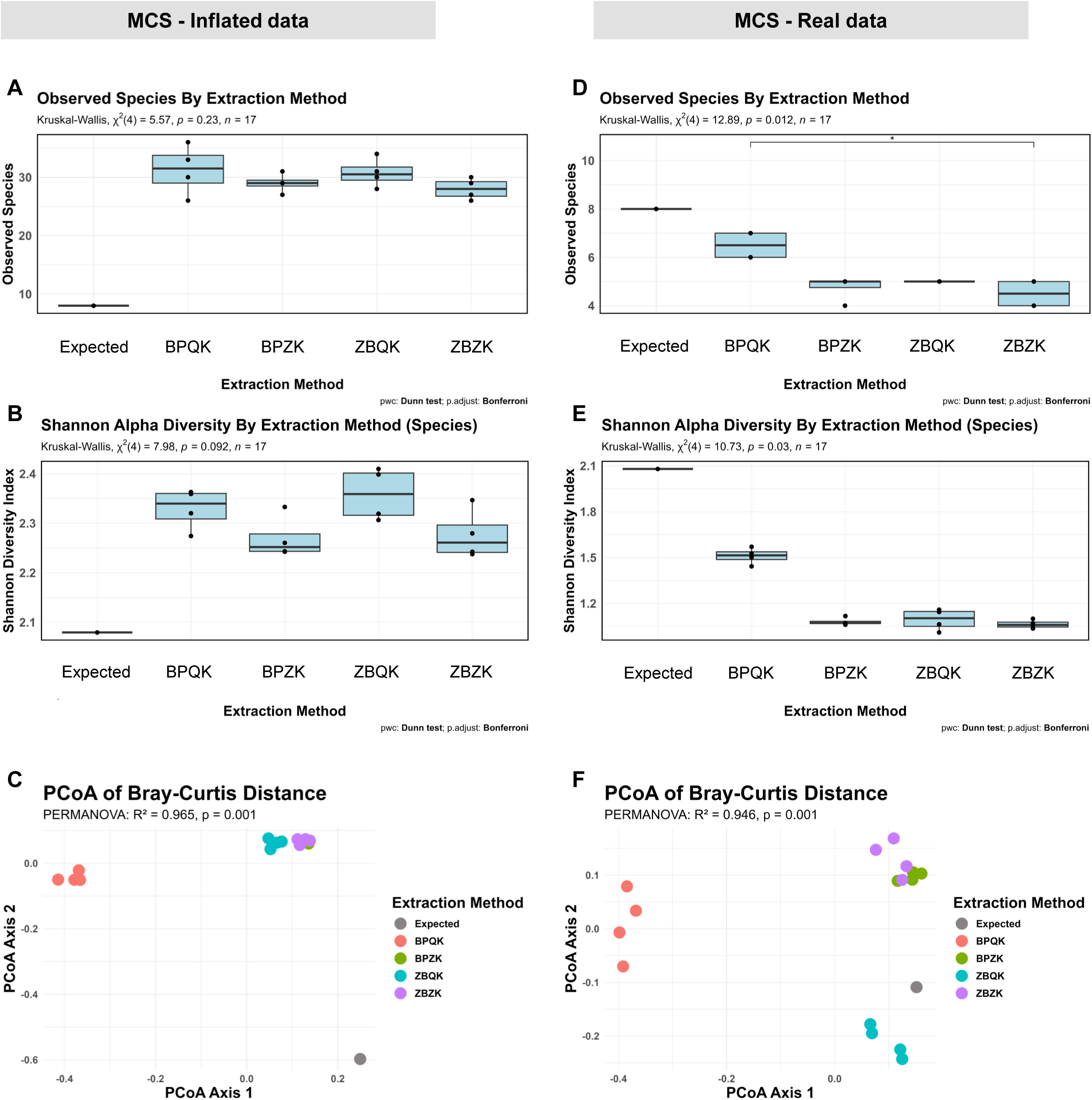
Comparison of alpha and beta diversity of the Microbial Community Standard (MCS) across four DNA extraction protocols before and after bioinformatic filtering. Panels **A–C** correspond to unfiltered data prior to removing misidentified taxa, and panels **D–F** correspond to the filtered dataset. (**A**) Observed species richness across protocols compared with the expected mock composition. (**B**) Shannon diversity index values calculated at the species level. (**C**) Principal Coordinates Analysis (PCoA) of Bray–Curtis dissimilarities displaying sample distribution by extraction method. (**D**) Observed species richness across protocols after filtering. (**E**) Shannon diversity index values of the filtered dataset calculated at the species level. (**F**) PCoA of Bray–Curtis dissimilarities displaying community distribution among extraction methods after filtering.

After filtering the MCS dataset at the species level, the observed species richness and Shannon diversity decrease across all protocols (**Figure 3D–3E**). Alpha diversity differed among extraction methods (Observed species: Kruskal–Wallis χ² = 12.89, p = 0.012; Shannon index: χ² = 10.73, p = 0.03), with BPQK showing the highest values. Beta diversity based on Bray–Curtis dissimilarities showed clear clustering by extraction method; BPQK formed a distinct cluster, whereas BPZK, ZBZK and ZBQK plotted closest to the expected composition (**Figure 3F**). PERMANOVA confirmed significant compositional differences among groups (R² = 0.946, p = 0.001). To ensure that these results were not driven by within-group dispersion, a test for homogeneity of multivariate dispersion was conducted. Distances to group centroids did not differ significantly among extraction methods for the MCS dataset (**Supplementary Figure 3A**), indicating comparable levels of variance across groups.

### *Drosophila* samples processed with the BPQK protocol had the most optimal genomic DNA extraction

Next, we evaluated the performance of the four extraction protocols in recovering bacterial gDNA from *D. melanogaster* gut samples, used here as a model animal tissue commonly employed to study the effects of experimental interventions on gut microbiota composition. These gut bacterial samples were processed in parallel with the MCS samples described above.

We first compared species composition across protocols (**Figure 4A**). Taxonomic classification revealed bacteria genera such as *Lactobacillus* and *Acetobacter* including species previously reported as part of the *Drosophila* commensal microbiota (reviewed in [^22–24^]), such as *Lactiplantibacillus plantarum* (formerly *Lactobacillus plantarum*)*, Levilactobacillus brevis* (previously known as *Lactobacillus brevis),* and *Acetobacter tropicalis.* Our data suggest that our Canton-S wild-type fly line also contained additional *Lactobacillus* and *Acetobacter* species, including *A. persici*, *Lactiplantibacillus paraplantarum*, *Lactiplantibacillus pentosus*, and *Lacticaseibacillus paracasei,* which were detected at >65 reads per sample. The relative abundance of these species varied among extraction protocols, with some samples also containing the anaerobic Gram-negative gammaproteobacterium *Frischella perrara*, originally described in the honeybee *Apis mellifera*[^25^] (**Figure 4A**). Replicate samples within each protocol showed generally consistent community profiles, though BPQK and BPZK displayed greater variability in the relative abundance of *Acetobacter* and *Lactobacillus* species. A small fraction of reads (<5%) were classified as “unknown” or represented low-abundance taxa (<65 reads).

**Figure 4.**
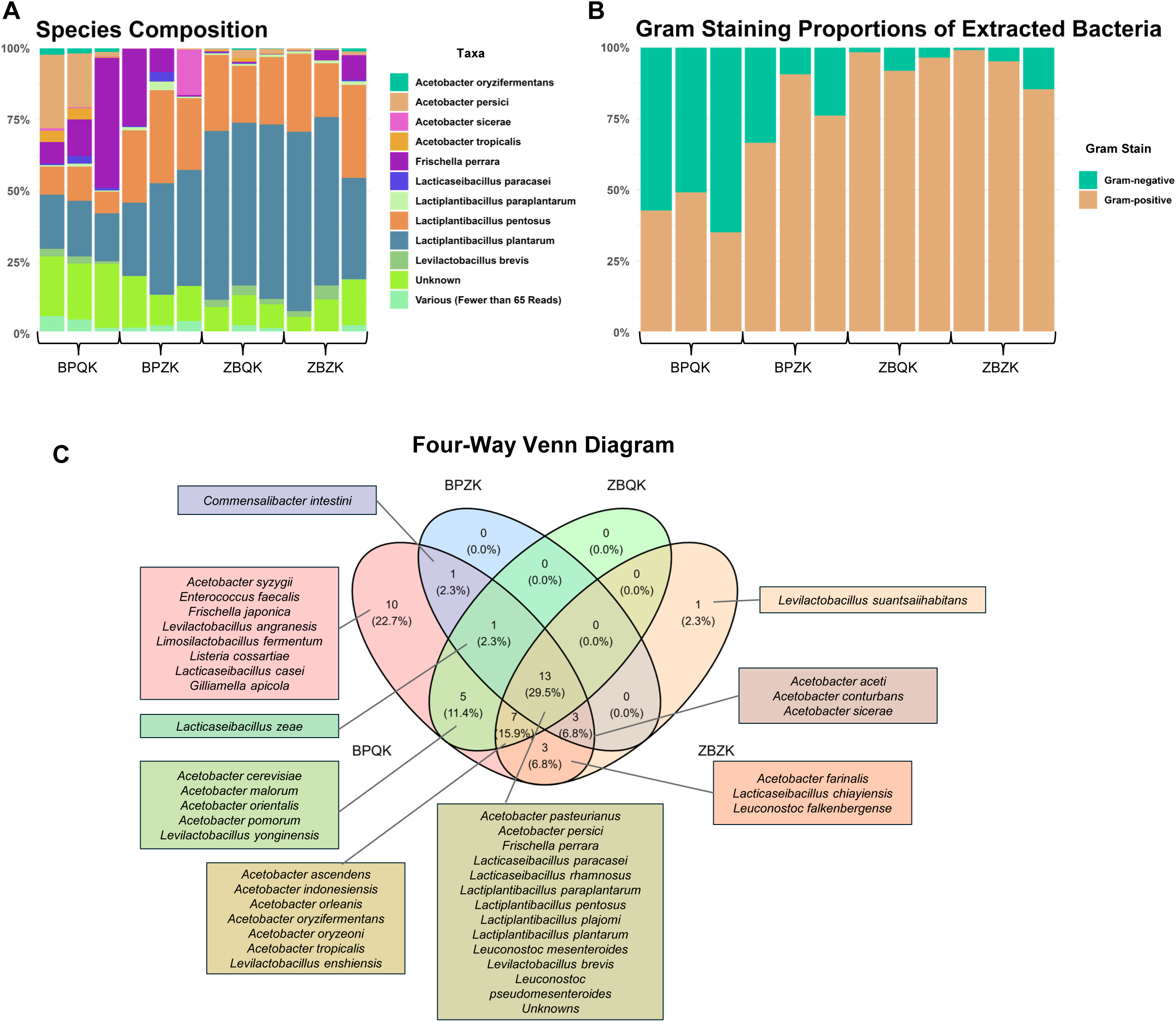
Taxonomic composition and Gram-stain distribution of *Drosophila melanogaster* gut microbiome samples across four DNA extraction protocols. **(A)** Relative abundance of bacterial species identified in gut samples processed using each extraction protocol. **(B)** Proportion of Gram-positive and Gram-negative bacterial species detected per protocol. **(C)** Four-way Venn diagram showing the number of shared and unique bacterial species identified across protocols. Twelve bacterial species were common to all four methods including *F. perrara* and *L. plantarum.* The BPQK protocol was the only one to detect nearly all taxa except *Levilactobacillus suantsaiihabitans*, which appeared exclusively in the ZBZK protocol.

To confirm reliability of the taxonomic assignments, p-distances among reference sequences were calculated in MEGA12, and the relationships among detected species were visualized using Neighbor-Joining phylogenetic tree (**Supplementary Figure 4**).

Gram-staining classification of the identified taxa indicated that the BPQK protocol extracted both Gram-positive and Gram-negative bacteria in nearly equal proportions (**Figure 4B**). The BPZK protocol also recovered representatives of both groups, though Gram-positive species were slightly more abundant. In contrast, the ZBQK and ZBZK protocols extracted fewer Gram-negative taxa, showing a clear bias toward Gram-positive bacteria.

To compare overlap among detected taxa, a four way Venn diagram was constructed (**Figure 4C**). Twelve bacterial species (excluding unknowns) were shared among all protocols, and included *F. perrara* and *L. plantarum*. However, only the BPQK protocol successfully recovered gDNA for all taxa detected across methods, with the exception of *Levilactobacillus suantsaiihabitans*, which was uniquely identified in the ZBZK protocol. In addition, BPQK uniquely detected *Frischella japonica* and *Listeria cossartiae*, whereas ZBZK exclusively recovered *Levilactobacillus suantsaiihabitans*.

### *Drosophila* gut microbiota samples extracted with the BPQK protocol tended to have the highest alpha diversity

We analyzed alpha and beta diversity metrics from the gut bacteria datasets obtained across the four extraction protocols (**Figure 5**). Among all protocols, BPQK samples displayed the highest observed species richness, with a median of approximately 33 species (Kruskal–Wallis, χ²(3) = 7.86, p = 0.049; Bonferroni-corrected post-hoc tests significant), whereas the remaining protocols showed medians below 20 (**Figure 5A**). Consistent with these findings, Shannon diversity indices showed a higher, though not significant, trend for BPQK compared to the other protocols (**Figure 5B**).

**Figure 5.**
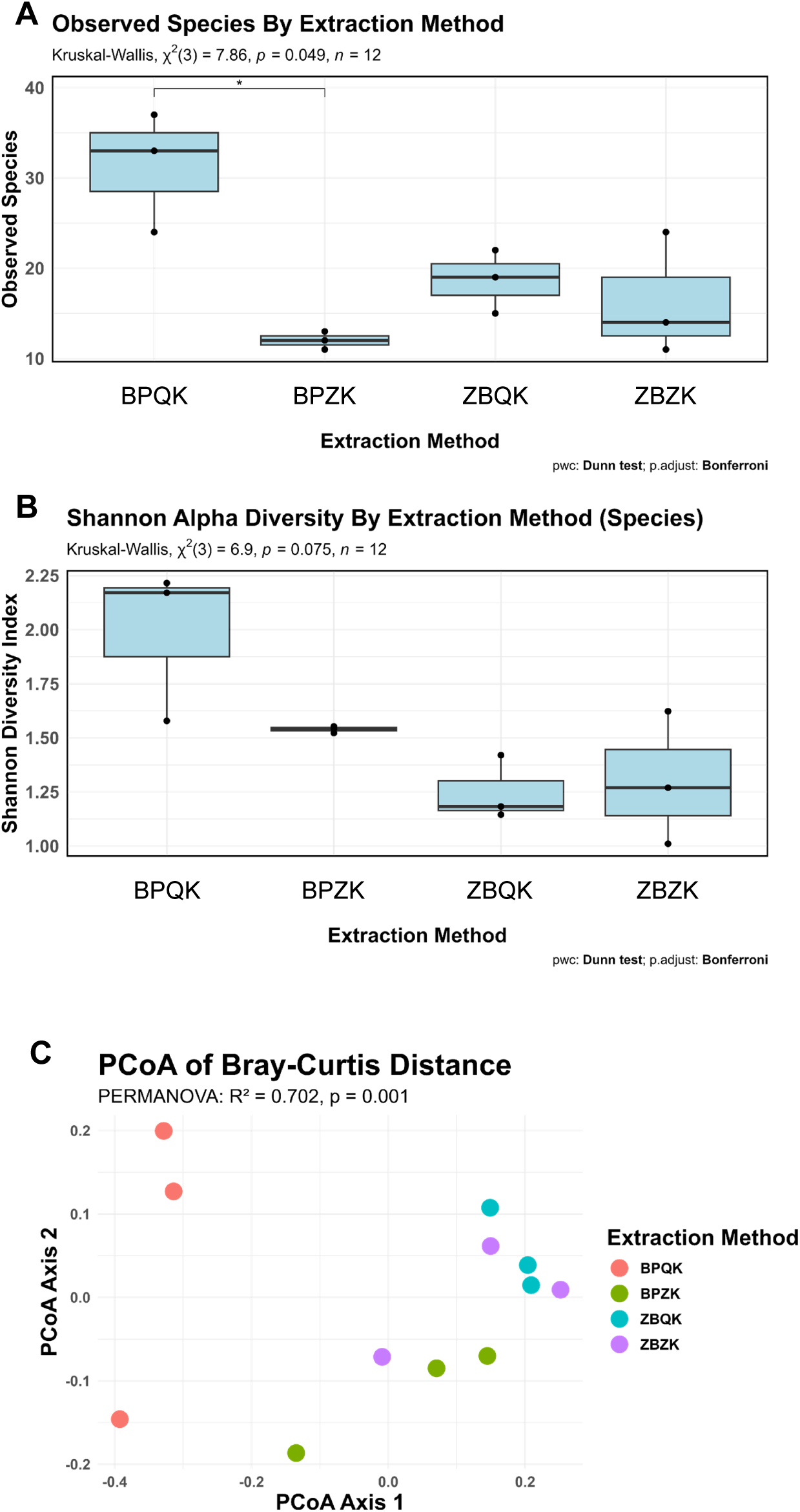
Alpha and beta diversity of *Drosophila melanogaster* gut microbiome samples across DNA extraction protocols. **(A)** Observed species richness across extraction protocols. **(B)** Shannon diversity index values for each protocol. **(C)** Principal Coordinates Analysis (PCoA) of Bray–Curtis dissimilarities showing sample clustering by extraction protocol.

Principal Coordinates Analysis (PCoA) based on Bray-Curtis dissimilarities revealed distinct clustering patterns by extraction method (**Figure 5C**). PERMANOVA confirmed a significant effect of extraction method on community composition (R² = 0.702, p = 0.001) with BPQK samples forming a cluster distinct from the other protocols. To ensure these results were not driven by unequal dispersion, we tested the homogeneity of multivariate dispersion; distances to group centroids did not differ significantly among methods for the *Drosophila* gut dataset (ANOVA: F= 0.906, p = 0.4796; **Supplementary Figure 3B**), indicating comparable variance within groups.

## Discussion

This study evaluated various DNA extraction methods and tissue lysis protocols to determine their effectiveness in profiling the gut microbiome of *D. melanogaster,* where an a MCS was included as a control. Our overarching goal was to develop a robust and reproducible workflow that combines commercially available extraction kits with a user-friendly bioinformatic pipeline, facilitating implementation in laboratories with diverse technical and computational resources. Results demonstrate that the choice of extraction method significantly influences microbial diversity and composition in both *D. melanogaster* tissue samples and an MCS, with distinct impacts on species detection, alpha diversity, and beta diversity. Importantly, our findings reveal protocol-specific biases in the recovery of Gram-positive versus Gram-negative bacteria, with the BPQK protocol yielding the most balanced taxonomic profile in both MCS, and more notably, the *Drosophila* gut samples. Similar biases in estimates of communities due to extraction protocols has been reported for human stool samples[^18^] or synthetic communities of human-associated bacteria[^19^].

The composition of the MCS revealed discrepancies between the expected and observed microbial profiles. The detection of additional genera not included in the MCS could reflect either potential contamination, such as the ‘kitome’ (bacteria coming from the kits), sequencing artifacts, or assignment misclassifications[^20^]. To control for contamination, we sequenced a non-template control (NTC), which consisted of adding sterile, nuclease-free water to the first PCR reaction instead of gDNA; this control was processed alongside all test samples throughout the sequencing process. The NTC contained only two low-abundance bacterial taxa, *Cutibacterium acnes* (formerly *Propionibacterium acnes*), a common human-associated bacterium, and a single environmental taxon (*Roseateles saccharophilus*), neither of which was detected in the MCS or *Drosophila* gut samples. Moreover, we did not detect any taxa entirely unexpected for the MCS or *Drosophila* datasets, such as typical human skin or environmental contaminants. This suggests that, in our case, the kits, pestles, and beads did not introduce additional contamination. However, a full kitome control, which would include reagent-only extractions processed in parallel with the samples, remains warranted for future studies to comprehensively rule out reagent-derived contaminants.

To further assess the contribution of sequencing errors, we analyzed the false-positive taxa detected in the inflated MCS dataset using phylogenetic reconstruction. Most false positives clustered within the same genera as their corresponding true positives, with pairwise sequence identities exceeding 99% in most cases. This high sequence similarity suggests that these discrepancies likely result from basecalling inaccuracies associated with the ∼1-3% error rate reported for Oxford Nanopore sequencing[^26,27^], rather than from true species-level differences. These findings emphasize the need for caution when interpreting species-level classifications from full-length 16S rRNA data and highlight the value of confirmatory sequencing using complementary platforms.

Interestingly, even the BPQK method, which performed best by detecting 7 out of the 8 species expected in the MCS, was not able to detect *Listeria monocytogenes* at the species level, although *Listeria* was identified at the genus level. This Gram-positive bacterium is known for its thick, teichoic-acid-rich peptidoglycan cell wall[^28^] but should, in principle, be easier to lyse than the spore-forming *Bacillus subtilis*[^29^], which was successfully detected by all protocols. This suggests that *L. monocytogenes* sequences may cluster with other *Listeria* species due to their high 16S rRNA gene sequence similarity. Indeed, in the inflated MCS dataset, at least nine *Listeria* species were identified, with *Listeria cossartiae* detected in all samples and showing the highest relative abundance among the genus. Phylogenetic analysis using MEGA12 confirmed that the 16S gene sequence of *L. cossartiae* shares >99% identity with *L. marthii, L. swaminathanii, L. innocua*, and *L. welshimeri*, and ∼98% with *L. monocytogenes* (BLASTn), supporting the likelihood of misassignment at the species level when using 16S data alone.

The absence of *E. coli,* together with *B. subtilis* in some protocols, despite *E. coli* being easily lysed[^29^], suggests that differences in DNA recovery may not stem solely from incomplete lysis but potentially from premature lysis and subsequent DNA degradation in more fragile species. Similarly, other studies have reported the absence of *E. coli* DNA in synthetic community samples where *E. coli* was known to be present[^19^]. Together, these findings underscore the importance of balancing enzymatic and mechanical disruption steps to avoid compromising genomic DNA integrity during extraction.

Species-level analysis of *Drosophila* gut samples further highlighted the variability among lysis/extraction protocols. The BPQK protocol consistently recovered *L. plantarum*, *A. percici*, and *F. perrara*, suggesting broad sensitivity for both Gram-negative and Gram-positive taxa. In contrast, BPZK showed high abundances of *L. plantarum* and *L. pentosus*, but reduced *Acetobacter* spp., while the Zymo-based protocols (ZBQK and ZBZK) primarily detected *L. plantarum* and *L. pentosus*, yet displayed a reduced detection of *F. perrara*. Together, these results demonstrate that the extraction method directly shapes the resulting taxonomic profile, with BPQK providing the most comprehensive representation of the fly gut community containing both Gram-negative (*Acetobacter*) and Gram-positive (*Lactiplantibacillus*) bacteria.

Alpha diversity measures also differed significantly among protocols. In the MCS, all protocols overestimated diversity relative to the expected, but after filtering, BPQK achieved the most accurate recovery of the true community. In the *Drosophila* gut samples, BPQK displayed the highest observed richness and Shannon diversity, capturing a broader range of taxa than the other methods. BPZK showed moderately high diversity but fewer observed species, while both Zymo-based protocols yielded lower and more uniform values. These findings suggest that, while Zymo kits perform well in mock standards, the Qiagen-based BPQK protocol is more effective for complex biological samples, such as fly gut tissue.

Beta diversity analyses reinforced these observations. In both datasets, Bray–Curtis dissimilarity revealed clear protocol-dependent clustering, with BPQK samples forming distinct groups. In the *Drosophila* gut samples, the BPQK again clustered separately, consistent with its broader taxonomic recovery and reproducible community structure.

The observed differences in microbial composition, richness, and clustering patterns underscore the critical role of DNA extraction methods in shaping microbiome data. Protocols that effectively lyse both Gram-positive and Gram-negative bacteria, such as BPQK, are essential for comprehensive microbial profiling. In contrast, the Gram-positive bias observed in Zymo-based protocols limits detection of key taxa such as *Acetobacter*. These protocol-specific differences highlight that even small variations in enzymatic or mechanical lysis can have substantial effects on apparent diversity and abundance. Together, the MCS and gut microbiome analyses demonstrate that discrepancies in detected taxa can arise from both the lysis/extraction method and sequencing artifacts, rather than from true biological variation among studies. Collectively, these findings underscore the importance of rigorous quality control, as reliable extraction and sequencing protocols should recover all expected taxa while minimizing false positives. To fully capture bacterial diversity within a sample, combining or splitting samples for complementary extraction methods may enhance the recovery of both Gram-positive and Gram-negative taxa.

### Limitations and Future Studies

Although this study provides valuable insights into the impact of DNA extraction methodology on microbiome composition using full-length 16S Nanopore sequencing, several limitations should be acknowledged. First, the number of biological samples analyzed was modest, which may limit the statistical power to detect subtle community differences. However, this work prioritized methodological validation over population-level inference, and the consistent patterns observed across replicates support the reliability of the findings.

Second, while Oxford Nanopore Technologies (ONT) enables accessible full-length 16S sequencing, results were not cross-validated with a second sequencing platform such as PacBio HiFi. Future studies could strengthen taxonomic resolution and error correction by integrating hybrid sequencing approaches or by submitting extracted gDNA for parallel sequencing using alternative technologies. Although full-length 16S sequencing provides improved resolution, species-level classifications should still be interpreted cautiously, and confirmatory sequencing using alternative platforms is recommended.

Third, the ONT 16S Barcoding Kit used in this study provides only 24 unique barcodes, constraining throughput and necessitating two sequencing runs. This introduces potential batch effects; however, identical reagents, flow cell chemistry, and analysis parameters were applied across runs to minimize technical variation. Additionally, specific samples were sequenced in both runs as internal controls to assess consistency and detect potential run-to-run variability.

Fourth, primer design may also influence taxonomic representation. The universal 27F primer used in this study contained a “CC” at the degenerate positions, whereas alternative versions incorporating “YM” or “CM” degeneracies have been shown to improve amplification across diverse bacterial taxa. Such primer sequence differences could affect the detection of certain lineages, particularly those with mismatches at the 27F binding site. Future work could compare these primer variants to evaluate their impact on taxonomic coverage and relative abundance estimates.

Fifth, our taxonomic classification depended on the databases available within EPI2ME. Because our goal was to create a workflow accessible to laboratories with limited computing infrastructure, we used this cloud-based platform to avoid computationally intensive basecalling and alignment steps. However, the limited reference databases in EPI2ME resulted in several unclassified reads. This limitation can be mitigated by uploading custom-curated databases to EPI2ME, after which species abundance tables can be processed through the R pipeline provided in this study. This workflow provides a practical alternative to existing pipelines such as NanoCLUST[^30^], NanoRTax[^31^], and EMU[^32^], which require access to high-performance computing infrastructure or paid cloud-based resources.

Sixth, while this work focused on qualitative comparisons, future studies incorporating quantitative metrics, including DNA yield, integrity, and lysis efficiency, will help clarify the mechanisms driving extraction bias. Future optimization should also examine whether premature lysis compromises DNA integrity in fragile taxa, particularly in mixed communities containing both easily lysed and spore-forming bacteria. Similarly, Proteinase K incubation time may influence Gram-specific recovery; preliminary observations from our group suggest that shorter incubations (e.g., 3 hours) favor Gram-negative detection, whereas overnight incubations may bias toward Gram-positive bacteria. Additionally, further work should determine whether the high abundance of *Bacillus* observed in BPQK samples reflects genuine recovery of spore-forming bacteria or overrepresentation due to differential DNA retention or purification efficiency.

Despite these limitations, the experimental design successfully demonstrates clear protocol-dependent biases during DNA extraction and provides a framework for establishing standardized, reproducible workflows for Nanopore-based microbiome studies, particularly in low-resource or teaching laboratory environments.

### Conclusions

This study demonstrates that DNA extraction protocols exert a major influence on measures of microbial diversity and composition in *Drosophila melanogaster* gut microbiome studies. Among the approaches tested, Blue Pestle homogenization combined with the modified Qiagen DNeasy Blood & Tissue Kit (BPQK) provided the most balanced recovery of Gram-positive and Gram-negative taxa and yielded the highest species richness and diversity. Nonetheless, all protocols introduced method-specific biases that affected microbial profiles, emphasizing the need for careful method selection and validation in microbiome research. Moving forward, the development and adoption of standardized, user-friendly workflows, using commercially available kits and accessible bioinformatic tools, will be essential to enhance accuracy, reproducibility, and cross-study comparability in long-read microbiome analyses.

Protocols for full-length 16S rRNA sequencing should also be optimized for each tissue or sample type studied (e.g., *Drosophila* gut) and published in a standardized manner, similar to the approach of the Earth Microbiome Project, which established reference protocols for 16S amplicon sequencing across diverse sample types such as feces, oral, skin, soil, and water[^33,34^]. This is particularly important as ongoing efforts aim to develop a roadmap for establishing guidelines that promote equitable sequence data reuse and interoperability in public microbiome datasets, which should also include detailed documentation of the genomic DNA extraction protocols used[^35^].

## Materials & Methods

### *Drosophila melanogaster* Husbrandy and Gut Microbiome Sample Preparation

Wild-type *Drosophila melanogaster* Canton-S flies were raised in a 25L incubator with 12h:12h light:dark cycle with ∼60% humidity. Newly eclosed female flies were collected, mated and aged for five days, and then dissected. Prior to dissections, flies were surface-sterilized by washing in 70% ethanol for 30 s followed by two washes in sterile 1x phosphate-buffered saline (PBS) (Fisher Cat. No. BP399500). Guts were dissected under flame in drops of sterile 1x PBS on a sterile Petri dish. Crops and Malpighian tubules were removed from each gut before pooling in 500 µL of sterile PBS on ice. After dissections and pooling (10 guts per sample), PBS was removed, leaving only the tissues. During dissection, samples were maintained in a 1.5 mL Eppendorf tube cooling rack pre-chilled at −20°C to prevent thawing and were subsequently stored at −80°C until processing.

### Microbial Mock Community Standard

The microbial MCS, ZymoBIOMICS Microbial Community Standard (Cat. No. D6300, Zymo Research) was obtained from Zymo Research (Irvine, CA) and was included in the present study as a control. The MCS contains cells of five Gram-positive bacteria (i.e., *Lactobacillus fermentum, Enterococcus faecalis, Staphylococcus aureus, Listeria monocytogenes,* and *Bacillus subtilis*), three Gram-negative bacteria (i.e., *Pseudomonas aeruginosa, Escherichia coli,* and *Salmonella enterica*), and two yeast species (i.e., *Saccharomyces cerevisiae* and *Cryptococcus neoformans*). Each bacterial species is present in equal relative abundances (12%), while both yeast species are present in relative abundances of 2%.

### Tissue Homogenization and Bacterial Genomic DNA Extraction

Two distinct tissue lysis methods were tested: blue pestle homogenization and bead-beating. For blue pestle homogenization, frozen guts samples (stored at −80°C as described above) or microbial community standard (MCS) samples were thawed on ice and mixed with the lysis buffer provided in the respective extraction kit, following the manufacturer’s protocol. The samples were then homogenized using a hand-held homogenizer (VWR® Disposable Pellet Mixers and Cordless Motor, Cat. No. 47747-370; VWR International) for approximately 3 min. For gut samples, the homogenization time was determined empirically, ending when no large tissue fragments were visible by direct inspection.

For bead beating homogenization, gut or MCS samples were first combined with the lysis buffer from the indicated kit and transferred to ZymoBIOMICS™ BashingBeads Lysis Tubes (2mm). The tubes were processed in a Vortex Genie 2 (Model G-560, Scientific Industries Inc.) equipped with a Horizontal Microtube Holder (SI-H524, Scientific Industries Inc.) for 20 min at maximum speed (3,200 rpm).

Following homogenization, genomic DNA was extracted using two commercial kits: i) ZymoBIOMICS™ DNA Miniprep Kit (Cat. No. D4300, Zymo Research) and the ii) Qiagen DNeasy Blood & Tissue Kit (Cat. No. 69504, Qiagen). The ZymoBIOMICS™ DNA Miniprep Kit protocol was followed without modification. For the Qiagen kit, the Proteinase K digestion step was extended overnight at 56°C according to the procedure described by Bombin *et al.* (2020)^14^. The MCS samples were processed simultaneously with gut samples.

This experimental design resulted in four treatment groups: Blue Pestles + Qiagen Kit (BPQK), Zymo Beads + Qiagen Kit (ZBQK), Blue Pestles + Zymo Kit (BPZK), and Zymo Beads + Zymo Kit (ZBZK) (**Table 1**). Each protocol was applied to both the MCS (n = 4 replicates) and *D. melanogaster* gut samples (n = 3 replicates, 10 guts per replicate). Sequencing was performed in two independent runs, with BPQK and ZBQK processed in Run 1, and BPZK and ZBZK in Run 2.

### Full-Length 16S rRNA gene Amplification

Full-length 16s rRNA genes were amplified by PCR using LongAmp® Taq DNA Polymerase (Cat. No. M0533S, New England BioLabs), and the universal bacterial primers 27F (5′-AGA GTT TGA TCC TGG CTC AG-3′) (Integrated DNA Technologies; Cat. No. 51-01-19-06) and 1492R (5′-ACG GCT ACC TTG TTA CGA CTT-3′) (Integrated DNA Technologies; Cat. No. 51-01-19-07). These primers amplify approximately 1500 bp of the 16S rRNA gene, allowing full-length 16S rRNA sequencing.

PCR conditions consisted of an initial denaturation at 94°C for 2 min, followed by 34 cycles 30 s, 58°C for 30 s, and 68°C for 90 s, with a final extension of 68°C for 90 s. PCR products were verified by agarose gel electrophoresis and purified using the Gene Choice PCR Purification Kit (Cat. No. 96-302, Genesee Scientific), according to the manufacturer’s instructions DNA quantity and purity were assessed with a Nanodrop spectrophotometer.

A non-template control (NTC) containing nuclease-free sterile water instead of genomic DNA was included in each PCR run. The NTC was processed identically to experimental samples through all subsequent steps, including sequencing, to serve as a control for contamination.

### Library Preparation and 16s rRNA Sequencing

Library preparation was performed using the Nanopore 16S Barcoding Kit 24 V14 (Cat. No. SQK-16S114.24, Oxford Nanopore), following the manufacturer’s Rapid Sequencing DNA protocol (Version 16S_9199_v114_revE_11Dec2024, Oxford Nanopore). Due to the kit’s limitation of 24 unique barcodes, two sequencing runs were required. To assess and control for potential batch effects, aliquots of the same extracted DNA samples were included in both runs under distinct barcodes.

Barcodes were attached to amplicons by PCR according to the manufacturer’s instructions. Barcoded libraries were quantified using a Qubit fluorometer (Thermo Fisher Scientific) and pooled in equimolar ratios. The pooled libraries were purified with AMPure XP Beads (Beckman Coulter) using 80% ethanol prepared in nuclease-free water, and DNA was eluted in 10uL of nuclease-free water. Final library concentrations were measured using the Qubit fluorometer, and 50 femtomoles of DNA were loaded for sequencing.

Sequencing was carried out on a GridION platform (Oxford Nanopore Technologies using R10.4.1 MinION flow cells; Cat. No. FLO-MIN114, Oxford Nanopore) operated under the high-fidelity (high-accuracy) sequencing mode to minimize error rates.

### Taxonomic Classification and Diversity Calculations

Sequencing data were analyzed using the 16S workflow (v1.5.0) in Oxford Nanopore’s cloud-based EPI2ME platform (https://epi2me.nanoporetech.com/), which includes the following steps: high-accuracy basecalling (using Dorado version 7.8.3 for run 1 and Dorado version 7.9.8 for run 2), demultiplexing, barcode trimming, read filtering and alignment to reference database. Taxonomic classification was performed with Minimap2 (version 2.28-r1209) against the NCBI 16S and 18S rRNA database (ONT given name: ncbi_targeted_loci_16s_18s.fna). Only reads between 1200 and 1800 base pairs and with a minimum quality score of 18 were retained for downstream analysis.

For diversity calculations and data visualization, the species abundance table generated by EPI2ME (**Supplementary Data 1**) was exported and processed in R (version 4.4.2) and RStudio (v2025.09.0+387) and a custom analysis script. All analyses were performed on a Beelink SER5 MAX Mini PC equipped with an AMD Ryzen 7 6800U processor (8 cores, 16 threads, up to 4.7 GHz), 32 GB RAM, and a 500 GB PCIe 4.0 SSD running Windows 11 Pro.

Rarefaction was performed to standardize sequencing depth (962 sequences for MCS and 1,108 sequences for gut samples) (**Supplementary Figure 5**). Alpha diversity metrics (including observed species richness and Shannon diversity index) were computed with the vegan package (v2.7-1). After the Kruskal–Wallis test, pairwise group comparisons were performed using the Dunn test with Bonferroni-adjusted p-values. Beta diversity was assessed using Bray–Curtis dissimilarities, and significance among extraction methods was tested by PERMANOVA computed with the vegan package (v2.7-1). Phylogenetic tree construction and UniFrac distance calculations (data not shown) were performed using the phyloseq package (v1.50.0). Gram-stain assignments were generated with the AMR package (v3.0.0), and data visualization was completed with ggplot2 (v4.0.0). Venn diagrams were produced with ggvenn (v0.1.10).

### Evolutionary relationships of taxa

Neighbor-joining trees for both MCS and *Drosophila* gut bacteria samples were constructed in MEGA12 (v12.0.14)[^36^]. Briefly, the evolutionary history was inferred using the Neighbor-Joining method[^37^], and the robustness of the tree topology was assessed using bootstrap analysis with 1,000 replicates[^38^]. The optimal tree for the MCS dataset had a total branch length of 0.908, and the tree for the gut samples had a total branch length of 0.931 (**Supplementary Figure 2 and 4).** Evolutionary distances were calculated using the p-distance method[^39^] and are expressed as the number of base differences per site. Bootstrap support values (percentages) are shown next to each internal node. The analyses included 111 nucleotide sequences for the MCS and 82 sequences for the gut samples (**Supplementary Figure 2 and 4**). Ambiguous positions were handled using the pairwise deletion option, resulting in a final dataset comprising 1,652 positions for MCS and 1,613 positions for gut samples.

## Data availability

Raw sequencing data are available at the NCBI Sequence Archive (SRA) under the accession number [to be provided prior to publication].

## Code availability

The custom script for downstream analysis of full-length 16S rRNAseq data generated from EPI2ME abundance tables is available on GitHub (https://github.com/carlosqui12/DNA_Extractions_2025h). Processed abundance tables are provided in **Supplementary Data 1**.

## Acknowledgments

We thank Dr. Lorraine Rodríguez-Rivera (Inter American University of Puerto Rico, San Juan, and Puerto Rico Science, Technology & Research Trust Research Institute) and Luis Acevedo (Puerto Rico Science, Technology & Research Trust Research Institute) for assistance with setting up the ONT GridION runs and data management. We also thank Dr. Cristina Martínez-Benito and the Scientific Writing class (BIOL 4999-423) at the University of Puerto Rico, Río Piedras, for their help with manuscript preparation, copy editing, and feedback.

We acknowledge funding support from the NIH-funded Increasing Diversity in Genomics for the Next Generation (IDGeNe) program (1R25HG012702) for supporting J.L.B.-D.V and from the NSF-funded RaMP: Research and Mentoring for Postbaccalaureates in Biological Sciences at the University of Puerto Rico (RaMP-UP) (2216584) to C.L.Q.-S. Additional support was provided by start-up funds from the University of Puerto Rico, Río Piedras Campus, the NIH–NIGMS COBRE (5P20GM103642 and 5P30GM149367), and the Catalyzer Research Grant (#2023-00056) from the Puerto Rico Science, Technology & Research Trust, awarded to I.A.R.-F.

## Author contributions

C.L.Q.-S. and J.L.B.-D.V. contributed equally to this work; Conceptualization: C.L.Q.-S., J.L.B.-D.V., and I.A.R.-F.; Methodology: C.L.Q.-S., J.L.B.-D.V., and M.U.; Investigation: C.L.Q.-S., J.L.B.-D.V., and M.U.; Formal analysis: C.L.Q.-S., J.L.B.-D.V., and T.M.S.-R.; Writing – original draft: C.L.Q.-S. and J.L.B.-D.V.; Writing – review & editing: C.L.Q.-S., J.L.B.-D.V., T.M.S.-R., and I.A.R.-F.; Supervision: I.A.R.-F. and T.M.S.-R.; Funding acquisition: I.A.R.-F.; Project administration: I.A.R.-F.; Bioinformatic support: T.M.S.-R.

## Additional information

**Supplementary information** The online version contains supplementary material available at [add link later].

## References

1. Sekirov, I., Russell, S. L., Caetano, L., Antunes, M. & Finlay, B. B. Gut Microbiota in Health and Disease. 10.1152/physrev.00045.2009.-Gut (2010) doi:10.1152/physrev.00045.2009.-Gut.

2. Turnbaugh, P. J. et al. The Human Microbiome Project. Nature vol. 449 804–810 (2007).

3. Hou, K. et al. Microbiota in health and diseases. Signal Transduction and Targeted Therapy vol. 7 (2022).

4. Afzaal, M. et al. Human gut microbiota in health and disease: Unveiling the relationship. Frontiers in Microbiology vol. 13 (2022).

5. Allaband, C. et al. Microbiome 101: Studying, Analyzing, and Interpreting Gut Microbiome Data for Clinicians. Clinical Gastroenterology and Hepatology vol. 17 218–230 (2019).

6. Ranjan, R., Rani, A., Metwally, A., McGee, H. S. & Perkins, D. L. Analysis of the microbiome: Advantages of whole genome shotgun versus 16S amplicon sequencing. Biochem. Biophys. Res. Commun. 469, 967–977 (2016).

7. Ursell, L. K., Metcalf, J. L., Parfrey, L. W. & Knight, R. Defining the human microbiome. Nutr. Rev. 70, S38–S44 (2012).

8. Clarridge, J. E. Impact of 16S rRNA Gene Sequence Analysis for Identification of Bacteria on Clinical Microbiology and Infectious Diseases. Clin. Microbiol. Rev. 17, 840–862 (2004).

9. Johnson, J. S. et al. Evaluation of 16S rRNA gene sequencing for species and strain-level microbiome analysis. Nat. Commun. 10, 5029 (2019).

10. Aja-Macaya, P. et al. Nanopore full length 16S rRNA gene sequencing increases species resolution in bacterial biomarker discovery. Sci. Rep. 15, 26486 (2025).

11. Hu, T., Chitnis, N., Monos, D. & Dinh, A. Next-generation sequencing technologies: An overview. Hum. Immunol. 82, 801–811 (2021).

12. Wang, Y., Zhao, Y., Bollas, A., Wang, Y. & Au, K. F. Nanopore sequencing technology, bioinformatics and applications. Nat. Biotechnol. 39, 1348–1365 (2021).

13. Zheng, P. et al. Nanopore sequencing technology and its applications. MedComm 4, e316 (2023).

14. Szoboszlay, M. et al. Nanopore Is Preferable over Illumina for 16S Amplicon Sequencing of the Gut Microbiota When Species-Level Taxonomic Classification, Accurate Estimation of Richness, or Focus on Rare Taxa Is Required. Microorganisms 11, (2023).

15. Karst, S. M. et al. High-accuracy long-read amplicon sequences using unique molecular identifiers with Nanopore or PacBio sequencing. Nat. Methods 18, 165–169 (2021).

16. Butler, I. et al. Standardization of 16S rRNA gene sequencing using nanopore long read sequencing technology for clinical diagnosis of culture negative infections. Front. Cell. Infect. Microbiol. 15, (2025).

17. Sohrabi, M. et al. The yield and quality of cellular and bacterial DNA extracts from human oral rinse samples are variably affected by the cell lysis methodology. J. Microbiol. Methods 122, 64–72 (2016).

18. Costea, P. I. et al. Towards standards for human fecal sample processing in metagenomic studies. Nat. Biotechnol. 35, 1069–1076 (2017).

19. Yuan, S., Cohen, D. B., Ravel, J., Abdo, Z. & Forney, L. J. Evaluation of methods for the extraction and purification of DNA from the human microbiome. PLoS ONE 7, (2012).

20. Salter, S. J. et al. Reagent and laboratory contamination can critically impact sequence-based microbiome analyses. BMC Biol. 12, (2014).

21. Felczykowska, A., Krajewska, A., Zielińska, S. & Łoś, J. M. Sampling, metadata and DNA extraction - important steps in metagenomic studies. Acta Biochim. Pol. 62, 151–160 (2015).

22. Broderick, N. A. & Lemaitre, B. Gut-associated microbes of Drosophila melanogaster. Gut Microbes 3, 307–321 (2012).

23. Wong, C. N. A., Ng, P. & Douglas, A. E. LowLdiversity bacterial community in the gut of the fruitfly *Drosophila melanogaster*. Environ. Microbiol. 13, 1889–1900 (2011).

24. Ludington, W. B. & Ja, W. W. Drosophila as a model for the gut microbiome. PLOS Pathog. 16, e1008398 (2020).

25. Engel, P., Kwong, W. K. & Moran, N. A. Frischella perrara gen. nov., sp. nov., a gammaproteobacterium isolated from the gut of the honeybee, Apis mellifera. Int. J. Syst. Evol. Microbiol. 63, 3646–3651 (2013).

26. Tyler, A. D. et al. Evaluation of Oxford Nanopore’s MinION Sequencing Device for Microbial Whole Genome Sequencing Applications. Sci. Rep. 8, (2018).

27. Zhang, T. et al. The newest Oxford Nanopore R10.4.1 full-length 16S rRNA sequencing enables the accurate resolution of species-level microbial community profiling. Appl. Environ. Microbiol. 89, (2023).

28. Abachin, E. et al. Formation of D-alanyl-lipoteichoic acid is required for adhesion and virulence of Listeria monocytogenes. Mol. Microbiol. 43, 1–14 (2002).

29. Islam, M. S., Aryasomayajula, A. & Selvaganapathy, P. R. A review on macroscale and microscale cell lysis methods. Micromachines vol. 8 (2017).

30. Rodríguez-Pérez, H., Ciuffreda, L. & Flores, C. NanoCLUST: a species-level analysis of 16S rRNA nanopore sequencing data. Bioinformatics 37, 1600–1601 (2021).

31. Rodríguez-Pérez, H., Ciuffreda, L. & Flores, C. NanoRTax, a real-time pipeline for taxonomic and diversity analysis of nanopore 16S rRNA amplicon sequencing data. Comput. Struct. Biotechnol. J. 20, 5350–5354 (2022).

32. Curry, K. D. et al. Emu: species-level microbial community profiling of full-length 16S rRNA Oxford Nanopore sequencing data. Nat. Methods 19, 845–853 (2022).

33. Marotz, C. et al. DNA extraction for streamlined metagenomics of diverse environmental samples. BioTechniques 62, 290–293 (2017).

34. Thompson, L. R. et al. A communal catalogue reveals Earth’s multiscale microbial diversity. Nature 551, 457–463 (2017).

35. Hug, L. A. et al. A roadmap for equitable reuse of public microbiome data. Nature microbiology vol. 10 2384–2395 (2025).

36. Kumar, S. et al. MEGA12: Molecular Evolutionary Genetic Analysis Version 12 for Adaptive and Green Computing. Mol. Biol. Evol. 41, msae263 (2024).

37. Saitou, N. & Nei, M. The neighbor-joining method: a new method for reconstructing phylogenetic trees. Mol. Biol. Evol. 10.1093/oxfordjournals.molbev.a040454 (1987) doi:10.1093/oxfordjournals.molbev.a040454.

38. Felsenstein, J. CONFIDENCE LIMITS ON PHYLOGENIES: AN APPROACH USING THE BOOTSTRAP. Evolution 39, 783–791 (1985).

39. Nei, M. & Kumar, S. Molecular Evolution and Phylogenetics. (Oxford University Press, Incorporated, Cary, 2000).

