## Supplementary Material for "Optimizing Tissue Lysis and DNA Extraction Protocols to Enhance Bacterial Diversity Profiling in the *Drosophila melanogaster* Gut Microbiome"

*** Correspondence:**
Co-corresponding Authors

**The supplementary materials contain the following:**

**Supplementary Data 1 –** Species abundance tables for MCS-Inflated, MCS-Real, and *Drosophila* gut samples, provided as a separate Excel file with individual tabs.

**Supplementary Data 2 –** Observed species corresponding to Supplementary Figure 1, provided as a separate Excel file with individual tabs per extraction protocol.

**Supplementary Figures 1–5 –** Included in this PDF document below.

**Supplementary Figures**

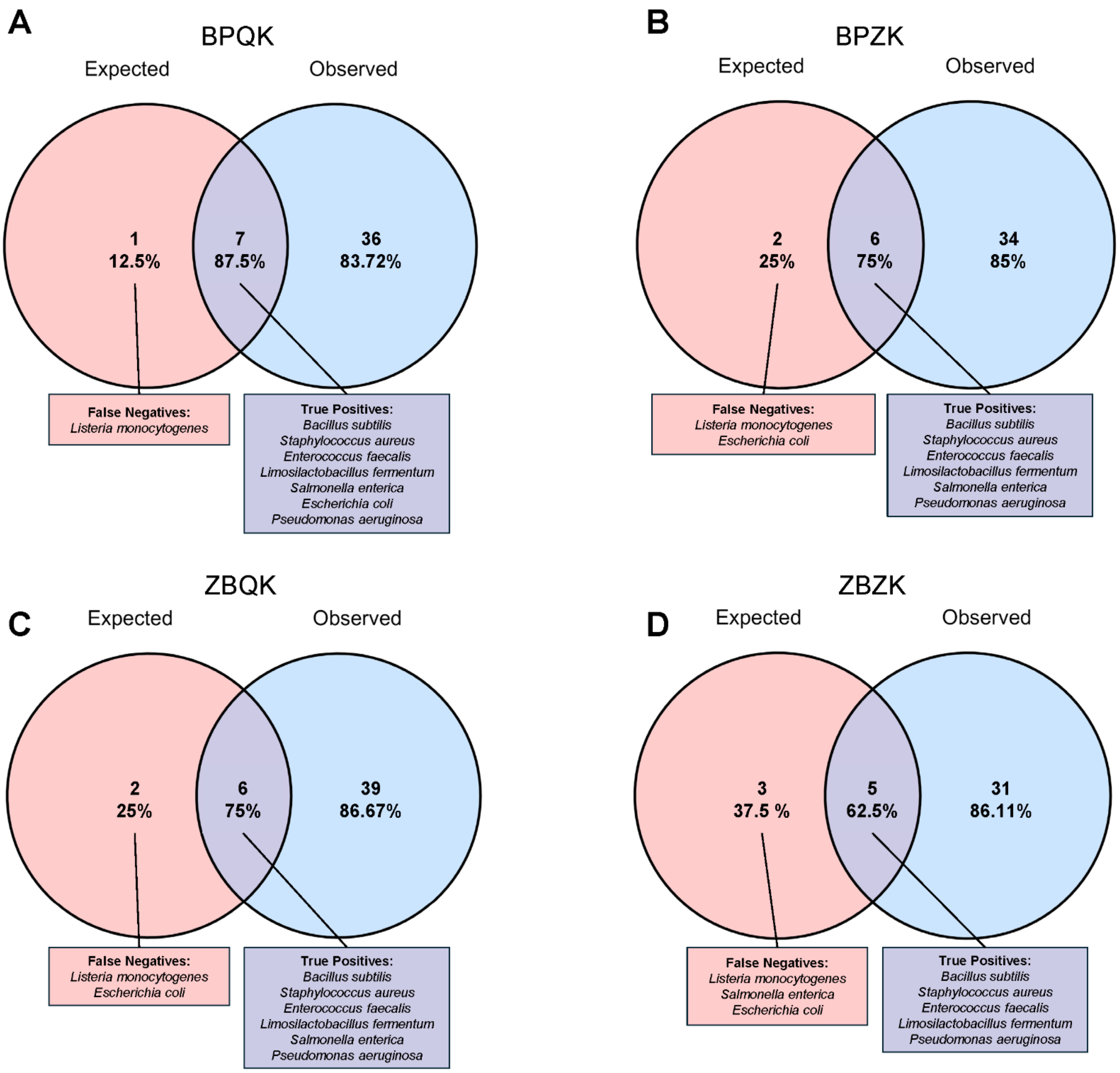


**Supplementary Figure 1 | Comparison of expected and observed bacterial species in the Microbial Community Standard (MCS) across DNA extraction protocols.** Venn diagrams show the overlap between expected and observed bacterial species detected after full-length 16S rRNA sequencing of the MCS using four extraction protocols. (**A)** BPQK (Blue Pestle + Qiagen Kit), (**B**) BPZK (Blue Pestle + Zymo Kit), (**C**) ZBQK (Bead Beating + Qiagen Kit), and (**D**) ZBZK (Bead Beating + Zymo Kit). Each diagram displays the proportion of true positives (species detected as expected) and false negatives (species expected but not detected). True-positive and false-negative taxa are listed for each protocol. BPQK recovered 7 of the 8 expected taxa, representing the highest detection rate among protocols. Refer to **Supplementary Data 2** for the complete list of observed species.

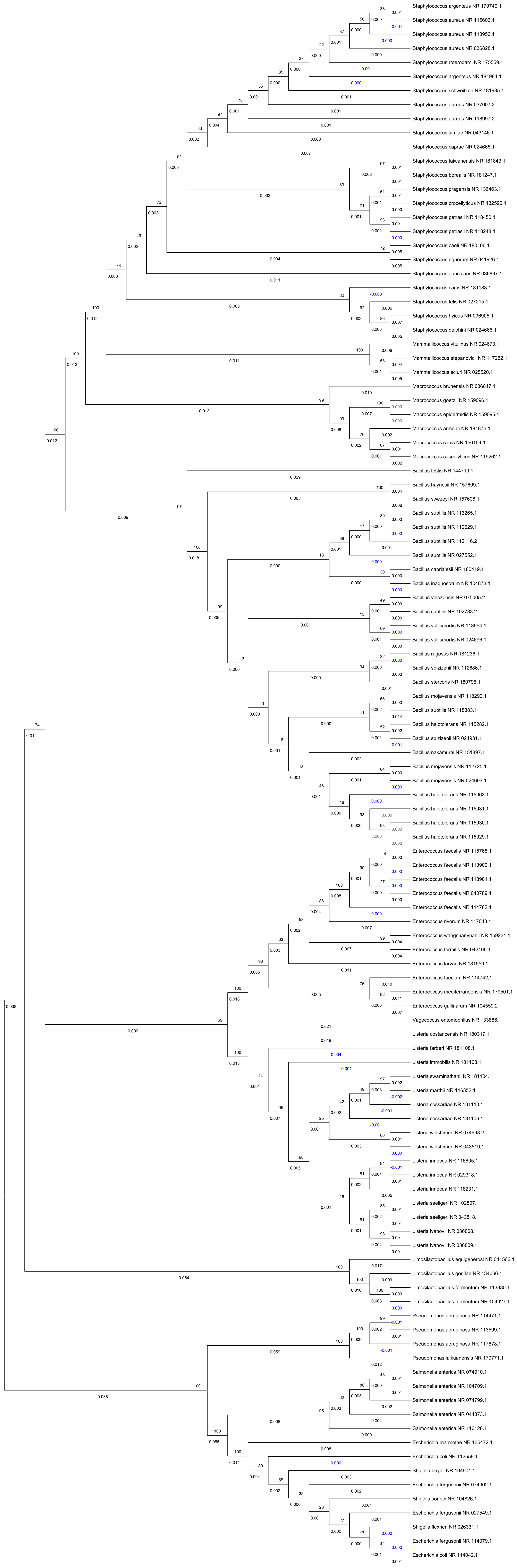
Staphylococcus argenteus NR 179740.1
Staphylococcus aureus NR 115606.1
Staphylococcus aureus NR 113956.1
Staphylococcus aureus NR 036828.1
Staphylococcus roterodami NR 175559.1
Staphylococcus argenteus NR 181984.1
Staphylococcus schweitzeri NR 181985.1
Staphylococcus aureus NR 037007.2
Staphylococcus aureus NR 118997.2
Staphylococcus simiae NR 043146.1
Staphylococcus caprae NR 024665.1
Staphylococcus taiwanensis NR 181843.1
Staphylococcus borealis NR 181247.1
Staphylococcus pragensis NR 136463.1
Staphylococcus croceilyticus NR 132590.1
Staphylococcus petrasii NR 118450.1
Staphylococcus petrasii NR 118248.1
Staphylococcus caeli NR 180106.1
Staphylococcus equorum NR 041926.1
Staphylococcus auricularis NR 036897.1
Staphylococcus canis NR 181183.1
Staphylococcus felis NR 027215.1
Staphylococcus hyicus NR 036905.1
Staphylococcus delphini NR 024666.1
Mammaliicoccus vitulinus NR 024670.1
Mammaliicoccus stepanovicii NR 117252.1
Mammaliicoccus sciuri NR 025520.1
Macrococcus brunensis NR 036847.1
Macrococcus goetzii NR 159096.1
Macrococcus epidermidis NR 159095.1
Macrococcus armenti NR 181876.1
Macrococcus canis NR 156154.1
Macrococcus caseolyticus NR 119262.1
Bacillus testis NR 144719.1
Bacillus haynesii NR 157609.1
Bacillus swezeyi NR 157608.1
Bacillus subtilis NR 113265.1
Bacillus subtilis NR 112629.1
Bacillus subtilis NR 112116.2
Bacillus subtilis NR 027552.1
Bacillus cabrialesii NR 180419.1
Bacillus inaquosorum NR 104873.1
Bacillus velezensis NR 075005.2
Bacillus subtilis NR 102783.2
Bacillus vallismortis NR 113994.1
Bacillus vallismortis NR 024696.1
Bacillus rugosus NR 181236.1
Bacillus spizizenii NR 112686.1
Bacillus stercoris NR 180796.1
Bacillus mojavensis NR 118290.1
Bacillus subtilis NR 118383.1
Bacillus halotolerans NR 115282.1
Bacillus spizizenii NR 024931.1
Bacillus nakamurai NR 151897.1
Bacillus mojavensis NR 112725.1
Bacillus mojavensis NR 024693.1
Bacillus halotolerans NR 115063.1
Bacillus halotolerans NR 115931.1
Bacillus halotolerans NR 115930.1
Bacillus halotolerans NR 115929.1
Enterococcus faecalis NR 115765.1
Enterococcus faecalis NR 113902.1
Enterococcus faecalis NR 113901.1
Enterococcus faecalis NR 040789.1
Enterococcus faecalis NR 114782.1
Enterococcus rivorum NR 117043.1
Enterococcus wangshanyuanii NR 159231.1
Enterococcus termitis NR 042406.1
Enterococcus larvae NR 181559.1
Enterococcus faecium NR 114742.1
Enterococcus mediterraneensis NR 179501.1
Enterococcus gallinarum NR 104559.2
Vagococcus entomophilus NR 133886.1
Listeria costaricensis NR 180317.1
Listeria farberi NR 181108.1
Listeria immobilis NR 181103.1
Listeria swaminathanii NR 181104.1
Listeria marthii NR 116352.1
Listeria cossartiae NR 181110.1
Listeria cossartiae NR 181106.1
Listeria welshimeri NR 074998.2
Listeria welshimeri NR 043519.1
Listeria innocua NR 116805.1
Listeria innocua NR 029318.1
Listeria innocua NR 118231.1
Listeria seeligeri NR 102807.1
Listeria seeligeri NR 043518.1
Listeria ivanovii NR 036808.1
Listeria ivanovii NR 036809.1
Limosilactobacillus equigenerosi NR 041566.1
Limosilactobacillus gorillae NR 134066.1
Limosilactobacillus fermentum NR 113335.1
Limosilactobacillus fermentum NR 104927.1
Pseudomonas aeruginosa NR 114471.1
Pseudomonas aeruginosa NR 113599.1
Pseudomonas aeruginosa NR 117678.1
Pseudomonas lalkuanensis NR 179771.1
Salmonella enterica NR 074910.1
Salmonella enterica NR 104709.1
Salmonella enterica NR 074799.1
Salmonella enterica NR 044373.1
Salmonella enterica NR 116126.1
Escherichia marmotae NR 136472.1
Escherichia coli NR 112558.1
Shigella boydii NR 104901.1
Escherichia fergusonii NR 074902.1
Shigella sonnei NR 104826.1
Escherichia fergusonii NR 027549.1
Shigella flexneri NR 026331.1
Escherichia fergusonii NR 114079.1
Escherichia coli NR 114042.1

**Supplementary Figure 2 | Phylogenetic relationships among bacterial taxa detected in the Microbial Community Standard (MCS).** Neighbor-Joining phylogenetic tree constructed in MEGA12 (v12.0.14) showing the evolutionary relationships among bacterial taxa identified in the MCS. Evolutionary distances were calculated using the p-distance method, and bootstrap support values (percentages) based on 1,000 replicates are shown next to each branch node. Branch lengths represent the number of base substitutions per site. The tree includes taxa representing both Gram-positive and Gram-negative bacteria recovered across the different DNA extraction protocols.

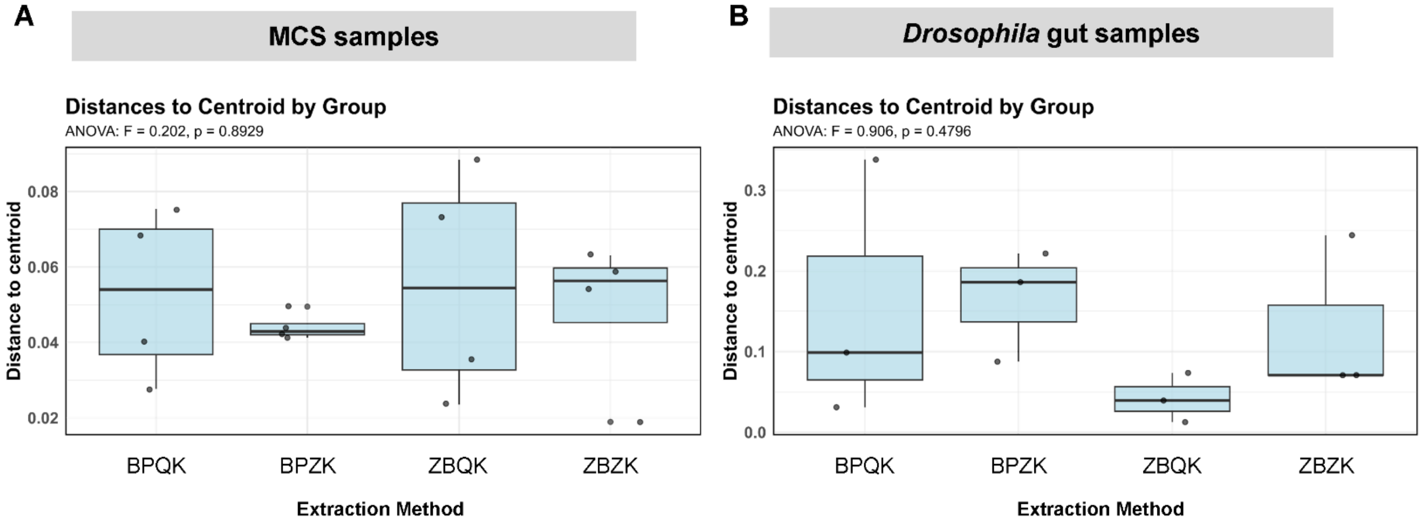


**Supplementary Figure 3**. **| Dispersion of microbial communities across extraction protocols.**

**(A)** Microbial Community Standard (MCS) samples and **(B)** *Drosophila melanogaster* gut samples showing distances to group centroids for each extraction protocol. Dispersion was evaluated using ANOVA to test for homogeneity of multivariate variance. No significant differences in dispersion were detected among extraction protocols for either dataset ($p > 0.05$).

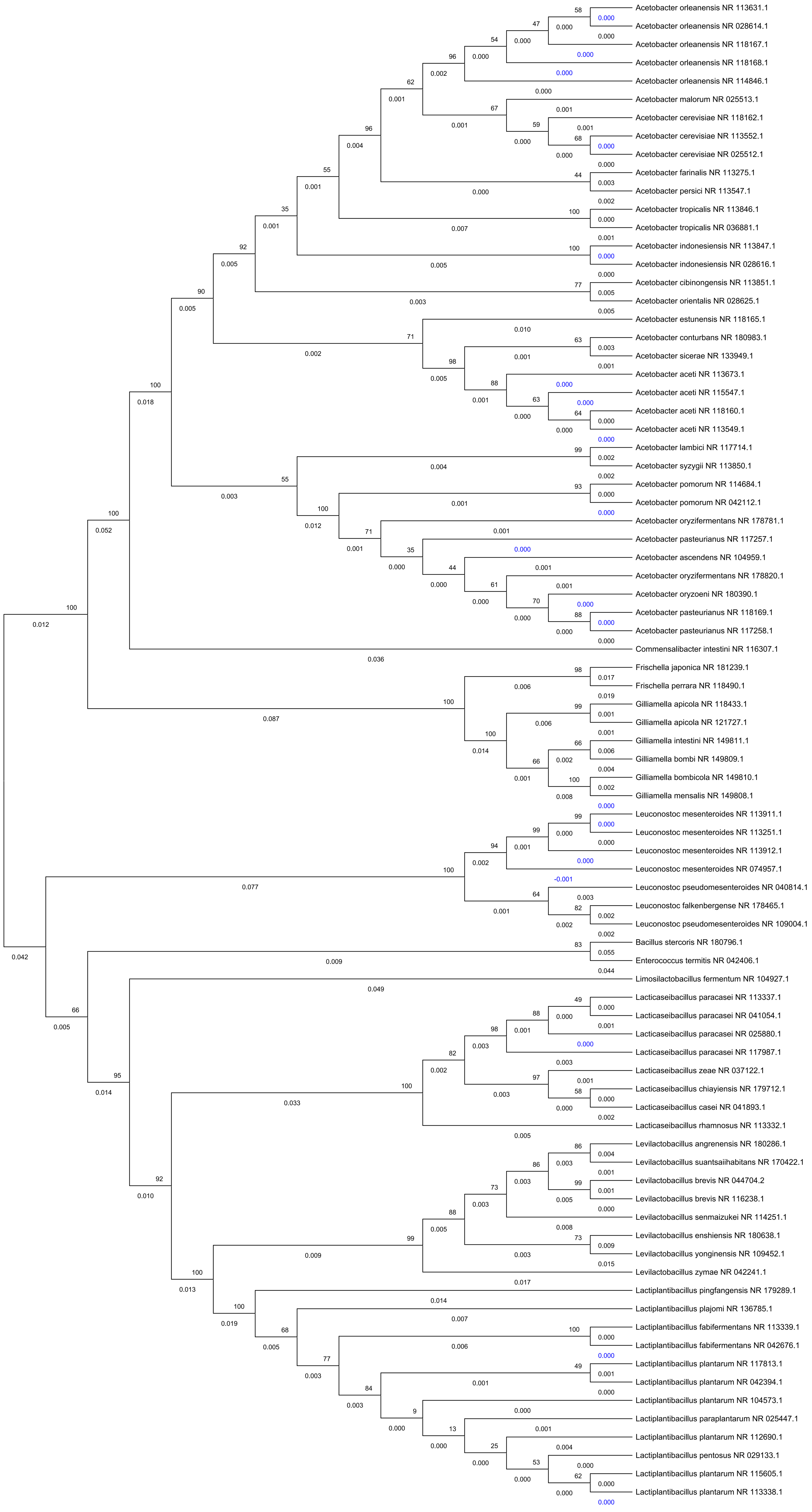
Acetobacter orleanensis NR 113631.1
Acetobacter orleanensis NR 028614.1
Acetobacter orleanensis NR 118167.1
Acetobacter orleanensis NR 118168.1
Acetobacter orleanensis NR 114846.1
Acetobacter malorum NR 025513.1
Acetobacter cerevisiae NR 118162.1
Acetobacter cerevisiae NR 113552.1
Acetobacter cerevisiae NR 025512.1
Acetobacter farinalis NR 113275.1
Acetobacter persici NR 113547.1
Acetobacter tropicalis NR 113846.1
Acetobacter tropicalis NR 036881.1
Acetobacter indonesiensis NR 113847.1
Acetobacter indonesiensis NR 028616.1
Acetobacter cibinongensis NR 113851.1
Acetobacter orientalis NR 028625.1
Acetobacter estunensis NR 118165.1
Acetobacter conturbans NR 180983.1
Acetobacter sicerae NR 133949.1
Acetobacter aceti NR 113673.1
Acetobacter aceti NR 115547.1
Acetobacter aceti NR 118160.1
Acetobacter aceti NR 113549.1
Acetobacter lambici NR 117714.1
Acetobacter syzygii NR 113850.1
Acetobacter pomorum NR 114684.1
Acetobacter pomorum NR 042112.1
Acetobacter oryzifermentans NR 178781.1
Acetobacter pasteurianus NR 117257.1
Acetobacter ascendens NR 104959.1
Acetobacter oryzifermentans NR 178820.1
Acetobacter oryzoeni NR 180390.1
Acetobacter pasteurianus NR 118169.1
Acetobacter pasteurianus NR 117258.1
Commensalibacter intestini NR 116307.1
Frischella japonica NR 181239.1
Frischella perrara NR 118490.1
Gilliamella apicola NR 118433.1
Gilliamella apicola NR 121727.1
Gilliamella intestini NR 149811.1
Gilliamella bombi NR 149809.1
Gilliamella bombicola NR 149810.1
Gilliamella mensalis NR 149808.1
Leuconostoc mesenteroides NR 113911.1
Leuconostoc mesenteroides NR 113251.1
Leuconostoc mesenteroides NR 113912.1
Leuconostoc mesenteroides NR 074957.1
Leuconostoc pseudomesenteroides NR 040814.1
Leuconostoc falkenbergense NR 178465.1
Leuconostoc pseudomesenteroides NR 109004.1
Bacillus stercoris NR 180796.1
Enterococcus termitis NR 042406.1
Limosilactobacillus fermentum NR 104927.1
Lacticaseibacillus paracasei NR 113337.1
Lacticaseibacillus paracasei NR 041054.1
Lacticaseibacillus paracasei NR 025880.1
Lacticaseibacillus paracasei NR 117987.1
Lacticaseibacillus zeae NR 037122.1
Lacticaseibacillus chiayiensis NR 179712.1
Lacticaseibacillus casei NR 041893.1
Lacticaseibacillus rhamnosus NR 113332.1
Levilactobacillus angrenensis NR 180286.1
Levilactobacillus suantsaiihabitans NR 170422.1
Levilactobacillus brevis NR 044704.2
Levilactobacillus brevis NR 116238.1
Levilactobacillus senmaizukei NR 114251.1
Levilactobacillus enshiensis NR 180638.1
Levilactobacillus yonginensis NR 109452.1
Levilactobacillus zymae NR 042241.1
Lactiplantibacillus pingfangensis NR 179289.1
Lactiplantibacillus plajomi NR 136785.1
Lactiplantibacillus fabifermentans NR 113339.1
Lactiplantibacillus fabifermentans NR 042676.1
Lactiplantibacillus plantarum NR 117813.1
Lactiplantibacillus plantarum NR 042394.1
Lactiplantibacillus plantarum NR 104573.1
Lactiplantibacillus paraplantarum NR 025447.1
Lactiplantibacillus plantarum NR 112690.1
Lactiplantibacillus pentosus NR 029133.1
Lactiplantibacillus plantarum NR 115605.1
Lactiplantibacillus plantarum NR 113338.1

**Supplementary Figure 4 | Phylogenetic relationships among bacterial taxa detected in the *Drosophila melanogaster* gut samples.** Neighbor-Joining phylogenetic tree constructed in MEGA12 (v12.0.14) showing the evolutionary relationships among bacterial taxa identified in *Drosophila melanogaster* gut samples. Evolutionary distances were calculated using the p-distance method, and bootstrap support values (percentages) based on 1,000 replicates are shown next to each branch node. Branch lengths represent the number of base substitutions per site. The tree includes taxa representing both Gram-positive and Gram-negative bacteria recovered across the different DNA extraction protocols.

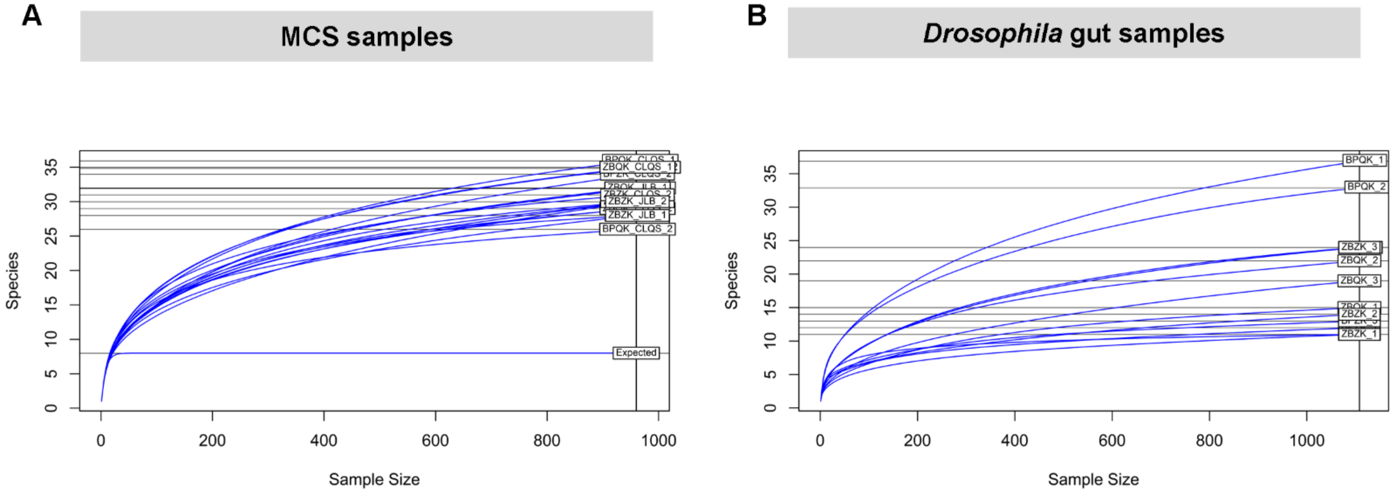


**Supplementary Figure 5 | Rarefaction curves of bacterial taxa detected across extraction protocols.**

**(A)** Rarefaction curves of the Microbial Community Standard (MCS) samples and **(B)** *Drosophila melanogaster* gut samples showing species accumulation as a function of sequencing depth. The curves illustrate the relationship between sample size and observed species richness for each extraction protocol, indicating that sequencing depth was sufficient to capture most taxa in each dataset. Rarefaction was based on 962 sequences for MCS samples and 1,108 sequences for gut samples.
